# Ototoxicity-related Changes in GABA Immunolabeling Within the Rat Inferior Colliculus

**DOI:** 10.1101/2024.07.29.603150

**Authors:** Avril Genene Holt, Ronald D. Griffith, Soo D. Lee, Mikiya Asako, Eric Buras, Selin Yalcinoglu, Richard A. Altschuler

## Abstract

Several studies suggest that hearing loss results in changes in the balance between inhibition and excitation in the inferior colliculus (IC). The IC is an integral nucleus within the auditory brainstem. The majority of ascending pathways from the lateral lemniscus (LL), superior olivary complex (SOC), and cochlear nucleus (CN) synapse in the IC before projecting to the thalamus and cortex. Many of these ascending projections provide inhibitory innervation to neurons within the IC. However, the nature and the distribution of this inhibitory input have only been partially elucidated in the rat.The inhibitory neurotransmitter, gamma aminobutyric acid (GABA), from the ventral nucleus of the lateral lemniscus (VNLL), provides the primary inhibitory input to the IC of the rat with GABA from other lemniscal and SOC nuclei providing lesser, but prominent innervation. There is evidence that hearing related conditions can result in dysfunction of IC neurons. These changes may be mediated in part by changes in GABA inputs to IC neurons. We have previously used gene micro-arrays in a study of deafness-related changes in gene expression in the IC and found significant changes in GAD as well as the GABA transporters and GABA receptors (Holt 2005). This is consistent with reports of age and trauma related changes in GABA (Bledsoe et al, 1995; Mossop et al, 2000; Salvi et al, 2000). Ototoxic lesions of the cochlea produced a permanent threshold shift. The number, intensity, and density of GABA positive axon terminals in the IC were compared in normal hearing and deafened rats. While the number of GABA immunolabeled puncta was only minimally different between groups, the intensity of labeling was significantly reduced. The ultrastructural localization and distribution of labeling was also examined. In deafened animals, the number of immuno gold particles was reduced by 78% in axodendritic and 82% in axosomatic GABAergic puncta. The affected puncta were primarily associated with small IC neurons. These results suggest that reduced inhibition to IC neurons contribute to the increased neuronal excitability observed in the IC following noise or drug induced hearing loss. Whether these deafness-diminished inhibitory inputs originate from intrinsic or extrinsic CNIC sources awaits further study.

## Introduction

The inferior colliculus (IC) is considered the integrative center of the auditory brainstem. The majority of ascending pathways from the cochlear nucleus (CN), lateral lemniscus (LL), and superior olivary complex (SOC), synapse in the IC before continuing to the medial geniculate body (MGB) and ultimately the cortex. The IC also integrates information from projection neurons located in the contralateral IC. Each of these ascending projections provide inhibitory afferent input to the neurons within the IC, the nature and the distribution of which have only been partially elucidated in the rat (Roberts and Ribak 1987; Adams and Mugnaini 1984; Oliver et al., 1994; Winer et al., 1995; and Merchan et al., 2005). Inhibitory circuits within the inferior colliculus shape the temporal integration of these ascending inputs. While the auditory pathways are reliant upon both gamma aminobutyric acid (GABA) and glycine for inhibitory neurotransmission, the inhibitory neurotransmitter GABA from the ventral nucleus of the lateral lemniscus provides the primary inhibitory input to the IC of the rat (Gonzalez-Hernandez et al., 1996) with GABA from other lemniscal, SOC nuclei, and the opposite IC providing a lesser, but nonetheless, prominent innervation of IC neurons. The distribution of these GABAergic inputs are described as both axosomatic and axodendritic with fewer GABA inputs apposing GABAergic IC neurons (Merchan et al., 2005, Oliver et al., 1994; Ito et al., 2009; Ito et al., 2018) than non-GABAergic IC neurons.

There is growing evidence that decreases in auditory input increase the excitability of IC neurons via a mechanism governed by GABAergic afferents. There have been several reports of activity dependent decreases in synaptic strength at GABAergic synapses in the IC (Vale and Sanes, 2002). Previously, we have used gene microarrays to study deafness related changes in gene expression in the IC and identified significant changes in the gene expression levels of GAD and GABA receptor subunits, as well as GABA transporters (Holt et al., 2005). These results are consistent with reports of age and noise related changes in GABA (Schatterman et al., 2008) as well as reports for a role of GABA in functional changes (Bledsoe et al., 1995; Helfert, et al., 1992; Mossop 2000; Salvi et al., 2000; Syka 2002; Vale and Sanes 2000; Vale et al., 2004; Szczepaniak and Moller 1995). Studies of the changes in GABAergic features in response to changes in activity are important for our future understanding of mechanisms underlying auditory system disorders such as presbyacusis, Meniere’s and tinnitus. Therefore, the principal purpose of this study is to determine the effect of diminished auditory activity on the morphology of IC neurons receiving GABAergic input as well as the number and distribution of GABAergic afferents across tonotopic regions of the central nucleus of the IC (CIC). Analysis of post-embedding immunocytochemistry for GABA at light and ultrastructural levels was used to assess changes following drug induced deafness.

## Materials and methods

### Animals

Ten adult female Sprague-Dawley rats (250-350g) purchased from Charles River Laboratories were used in this study in accordance with procedures approved by the University of Michigan Committee for the Use and Care of Animals. All animals were tested for normal hearing using auditory brainstem responses (ABRs) tested at frequencies of 2 kHz, 10 kHz, and 20 kHz, with sound pressure levels up to 100 dB SPL. Animals were randomly assigned either to the normal hearing group (n = 5) or the neomycin deafened group (n = 5).⍰ Fourteen days following the surgery, hearing was again tested before tissue harvest.

### Deafening Surgery

Rats were anesthetized with an i.m. injection of xylazine (8mg/kg) and ketamine (75mg/kg). To decrease pain and irritation of the wound following surgery, subcutaneous injections of 1% lidocaine-HCl were made locally prior to the first incision for both the right and the left ear. After exposing and opening the lateral wall of the bulla, 30 µl of 30% neomycin was infused into the cochlea through the round window over a 3 - 5 minute period, using a Hamilton syringe. Bilateral hearing threshold shifts of ≤ 10 dB or ≥ 65 dB for the normal hearing and deafened groups, respectively, were necessary to be included in the study. Hearing threshold requirements did not lead to exclusion of any animals from the study.

### Tissue Fixation and Processing

Fourteen days following surgery, animals were deeply anesthetized with 35% chloral hydrate and perfused transcardially, first with a Ca^+2^ free Ringer’s solution variant followed by a fixative comprising 1.25% glutaraldehyde and 2.0% freshly depolymerized paraformaldehyde. Following perfusion, brains were removed from the skulls and were subjected to a 90 minute post-fixation period in the same paraformaldehyde/glutaraldehyde solution. Tissue blocks containing the inferior colliculus were then cut into 150 μm sections using a vibratome (Vibratome 1000, Ted Pella, Inc.) and placed serially into four vials containing 0.12M NaPO_4_ buffer.

Sections were incubated for one hour in 1.0% osmium tetroxide and afterward counterstained with 2.0% uranyl acetate. Tissues were then dehydrated in a series of increasingly concentrated alcohols, culminating in a three-minute rinse in propylene oxide. Sections were then infiltrated in a 50:50 solution of propylene oxide:Epon812 overnight and subsequently embedded in pure Epon812 for four hours. Embedded sections were then flattened by compressing them between mylar sheets and glass plates and polymerizing them at 60° C for 48 hours.

To determine suitability for analysis, flattened sections were inspected under low magnification using a light microscope. Sections suitable for analysis were identified using anatomical landmarks consistent with those known to be present in sections through the mid rostro-caudal portion of the CIC (Figure 1). Following trimming of the blocks to the desired size, semi-thin sections (1 μm) were cut on a Reichert ultramicrotome using an 8 mm DDK diamond knife. Two adjacent sections were collected for use in GABA immunocytochemistry. Two different sections, one of which was collected just prior to the aforementioned sections and one collected just after, were immediately stained with Toluidine Blue for use in determining both the rostro-caudal position and subnuclei within the inferior colliculus. The next ten sections were discarded and then the next series of four sections was collected until 7 series of sections were collected (Series A-G).

**Figure 1.**
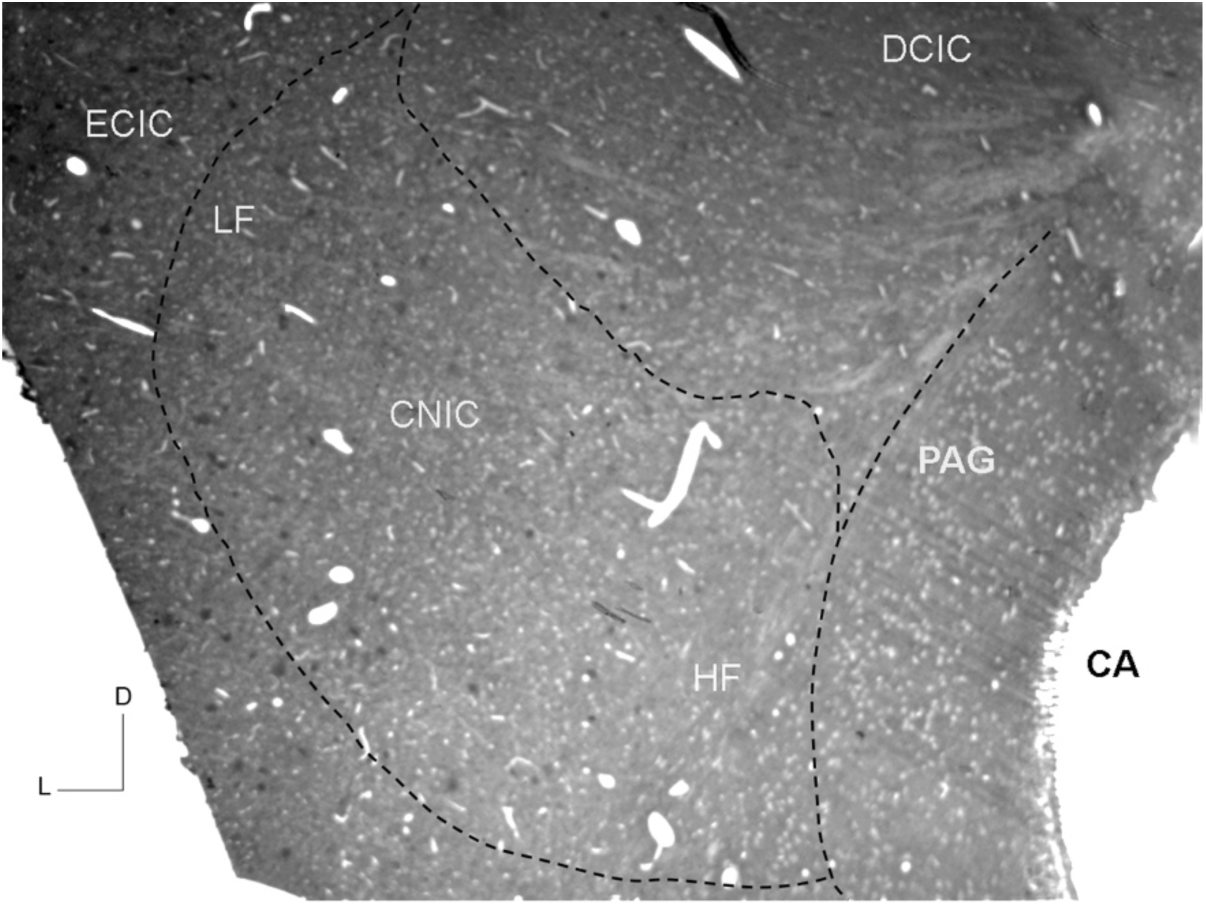
A plastic embedded section through the mid rostro-caudal inferior colliculus immunolabeled for GABA. At this level, three subdivisions of the inferior colliculus are visible: external cortex (ECIC), dorsal cortex (DCIC) and central nucleus (CNIC). PAG-periaqueductal gray; CA cerebral aqueduct; D - dorsal; L – lateral. LF=low frequency; HF=high frequency.

### Post-embedding immunocytochemistry for GABA

#### Light Microscopy

The 1 µm Epon812 embedded sections were etched with sodium ethoxide, rehydrated in decreasing concentrations of alcohol followed by water, and bleached in a 1.0% solution of sodium periodate (NaIO_4_). The adjacent sections were exposed to a 5% blocking solution for twenty minutes, and then incubated for 24-48 hours with 1:1000 rabbit anti-GABA (Strategic BioSolutions, Ramona, CA) antibody solution. Following incubation with the primary antisera, sections were incubated in a solution containing a biotinylated secondary antibody followed by incubation in ABC Vectastain Elite solution (Vector, Burlingame, CA). The DAB reaction was then carried out for 6 minutes.

#### Electron Microscopy

Once sections from the seventh series were collected, 70 – 80 nm sections (ultrathin sections) were collected for labeling and analysis. First, a 1 µm thin section was collected and stained for toluidene blue followed by ten ultrathin sections that were labeled for GABA using immunogold particles. Ultrathin sections were briefly etched with sodium ethoxide, and then incubated for 24 hours with rabbit anti-GABA (Calbiochem, San Diego, CA) antibody solution (1:250). Following incubation with the primary antisera, sections were incubated in a solution containing an immunogold conjugated secondary antibody (10 nm, Aurion EM Reagents Hatfield, PA).

### Analysis

#### Morphometry

Digital images from five sections from each of five rats in each group (25 sections) were acquired with a Spot camera (Diagnostic Instruments, Inc.) mounted onto a Ziess Axioplan microscope and analyzed using MetaMorph imaging software (Molecular Devices, Inc., Downingtown, Pennsylvania, USA). Nuclei for CIC neurons on average are no larger than the 12 µm interval between sections, this ensured that the same neuron would be not assessed twice. From each section analyzed, two areas of interest within both the high and low frequency regions of the central nucleus of the IC were selected which corresponded to the most ventromedial and dorsolateral aspects of the CIC, respectively (Figure 1). Stereology (MetaMorph imaging software; Molecular Devices, Inc.) was employed to identify and count neurons within each region. Within each selected area every neuron containing a nucleus, not touching another cell or touching the image edge was circled (Figure 2). Parameters such as area, perimeter, width, length, shape factor, and optical density were measured for each neuron.

**Figure 2.**
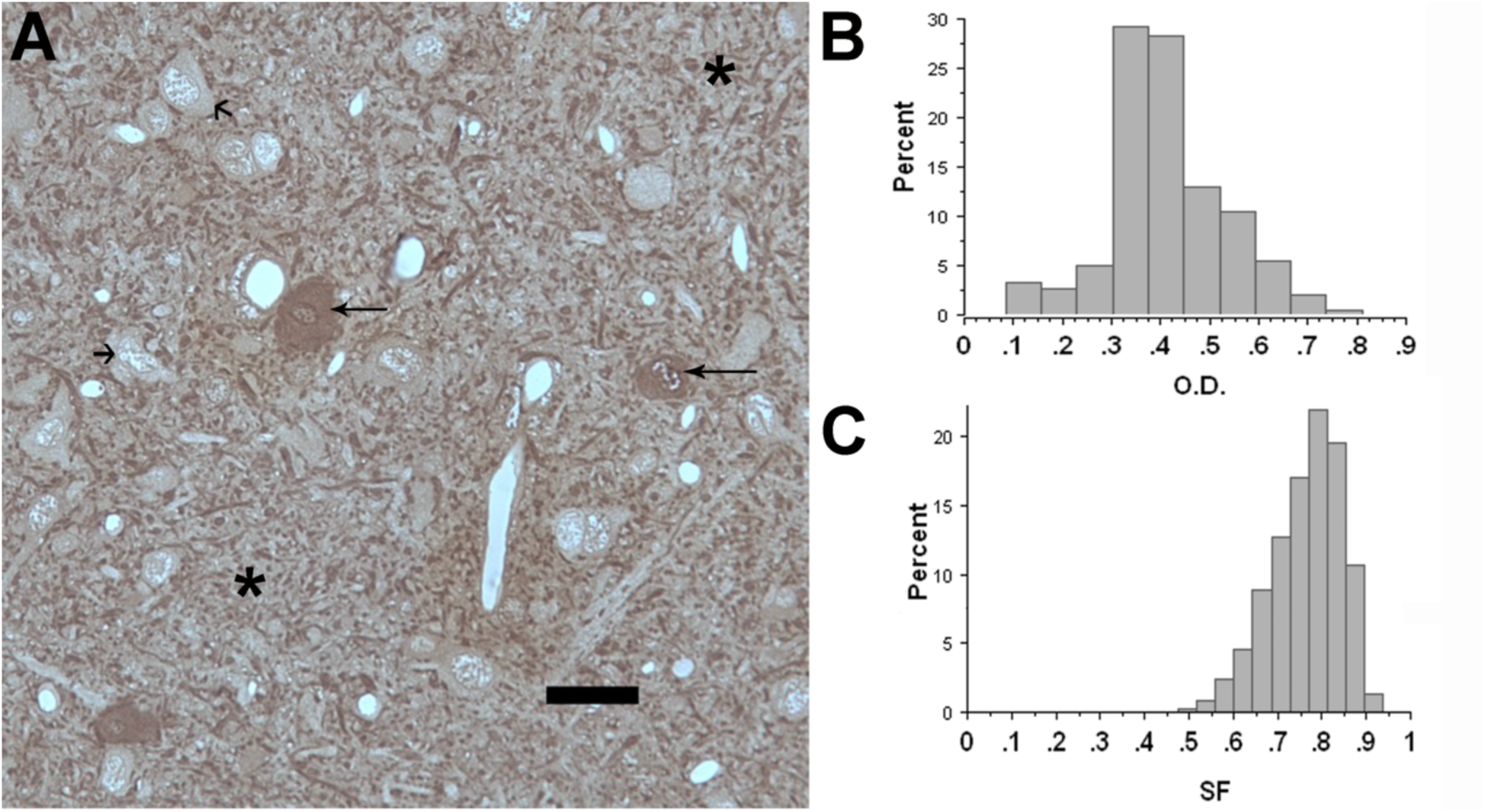
A photomicrograph of a high frequency reference area showing unlabeled and GABA labeled neurons as well as immunoreactive puncta (40x; A). Somas labeled for GABA (long arrows) and those that were unlabeled (short arrow) were present in each reference area. Asterisks denote the neuropil between somata. The percentage of somata as a function of optical density (O.D) are represented as a histogram (B). Somata roundness (SF) increases as the numbers approach one (x-axis; C). Optical density - O.D.; Shape factor – SF; Scale bar = 25 µm.

When using light microscopy, a neuron was characterized as being either labeled or unlabeled by assigning each cell in a particular image into one of two groups using a frequency distribution histogram based upon the cell’s optical density and then cells falling into the labeled group were further defined by intensity level (at least 5x over background). Additionally, GABA positive puncta were defined using size constraints (1.0 – 4.5 µm^2^) and shape (round to oval) criterion. For axosomatic labeling, the number of GABA immunoreactive puncta directly apposing each circled somatic profile was counted. For axodendritic labeling, GABA immunoreactive puncta in the neuropil was calculated as percent area using the defined size and shape constraints above, while excluding somata and axosomatic puncta (MetaMorph imaging software; Molecular Devices, Inc.). For ultrastructural analysis, the number of immuno-gold particles was counted in axosomatic and axodendritic boutons and compared in normal versus deafened animals. In order to be defined as an axosomatic bouton, the nucleus of the apposed soma needed to be clearly visible.

#### Statistical analyses

An analysis of variance (ANOVA) was performed (Statview, SAS Institute Inc., version 5.0). At the light level, comparisons between normal hearing and deafened groups were made for several parameters including, number of axosomatic puncta, cell area and shape for each cell type (labeled or unlabeled) from each of the 25 sections assessed in each group. For electron microscopy, comparisons were made for the number of immunogold particles present in either axosomatic or axodendritic boutons in the CIC of normal hearing versus deaf animals. For each parameter the averages were calculated per animal, with degrees of freedom equal to 1 in the numerator (deaf versus normal) and 8 in the denominator (five animals per group), with a confidence level of 95% necessary for significance. The Scheffe multiple comparison test was used for post-hoc comparisons.

## Results

### Light Microscopy

#### General morphology

Analysis of cell size, shape factor, and number of axosomatic and axodendritic puncta, as a function of either tonotopicity (high frequency versus low frequency) or staining density (labeled or unlabeled) demonstrated that darkly labeled GABAergic somata (10X over background) were the largest of all neurons (area = 158.11 µm^2^ ± 9.20 µm^2^; p<0.0001). On average, GABA immunoreactive somata (145.43 µm^2^ ± 6.53 µm^2^; p<0.0001) were larger than unlabeled somata (118.24 µm^2^ ± 1.86 µm^2^; p<0.0001). We also observed differences in shape factor when comparing GABA labeled versus unlabeled cells. The average somata that was unlabeled for GABA had larger shape factors (0.773 ± 0.03 vs 0.755 ± 0.03) and were therefore more round than those neurons labeled for GABA.

A gross assessment of immunocytochemistry for GABA in normal hearing subjects within the central nucleus of the IC revealed GABA labeled neurons throughout the nucleus (Figure 1) with the most labeled cells located laterally. While the total number of GABA positive neurons totaled fewer than one third of all CIC neurons counted, regardless of frequency region, a densely populated network of GABAergic puncta and dendrites could be fully appreciated throughout the neuropil (Figure 2A). Labeled cells ranged from lightly labeled to darkly labeled (optical density - O.D.; Figure 2B), and from round to fusiform in terms of shape (shape factor - SF; Figure 2C). Qualitative comparisons of GABA immunoreactive sections from normal hearing and deafened subjects showed a distinct difference in labeling (Figure 3). While GABA labeling (puncta and neurons) was robust in the normal group (Figure 3A, 3B), GABA labeling in the deafened group, while still detectable, was weak especially in the neuropil (Figure 3C, 3D, 3E). The shape of the GABA labeled somata were 5% more round following deafness (Figure 4C).

**Figure 3.**
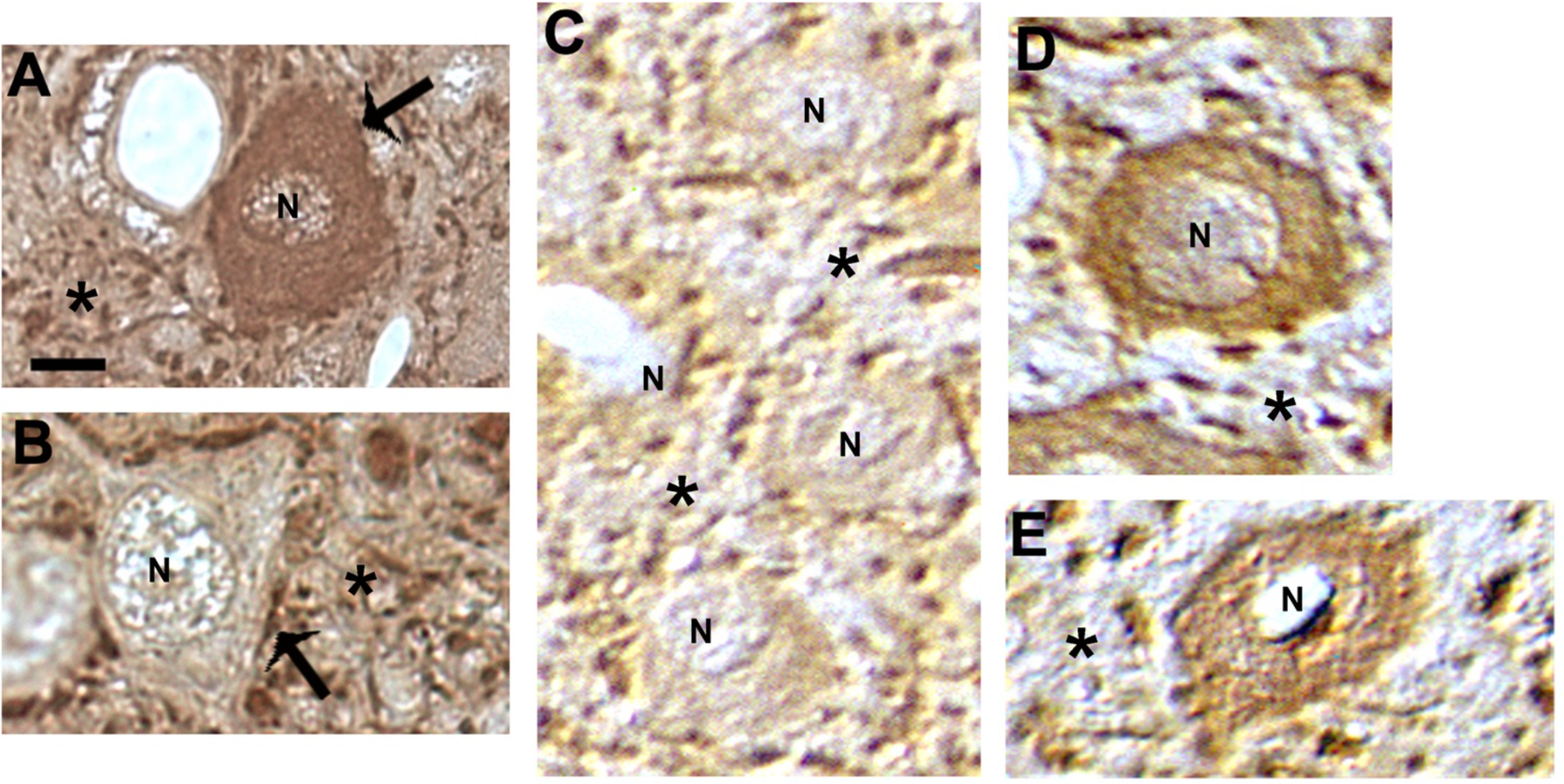
Light micrographs of somata with varying intensities of GABA immunolabeling. GABAergic puncta (indicated by arrows) are evident around both GABA labeled (A) and unlabeled somata (B). Numerous GABA labeled axosomatic and axodendritic puncta remain14 days after deafening (asterisks - *). However, the intensity of GABAergic puncta that are evident around the perimeter of the neuron and in the neuropil appears to decrease (C, D, E). N - Nuclei in all images; Scale bar = 10µm

**Figure 4.**
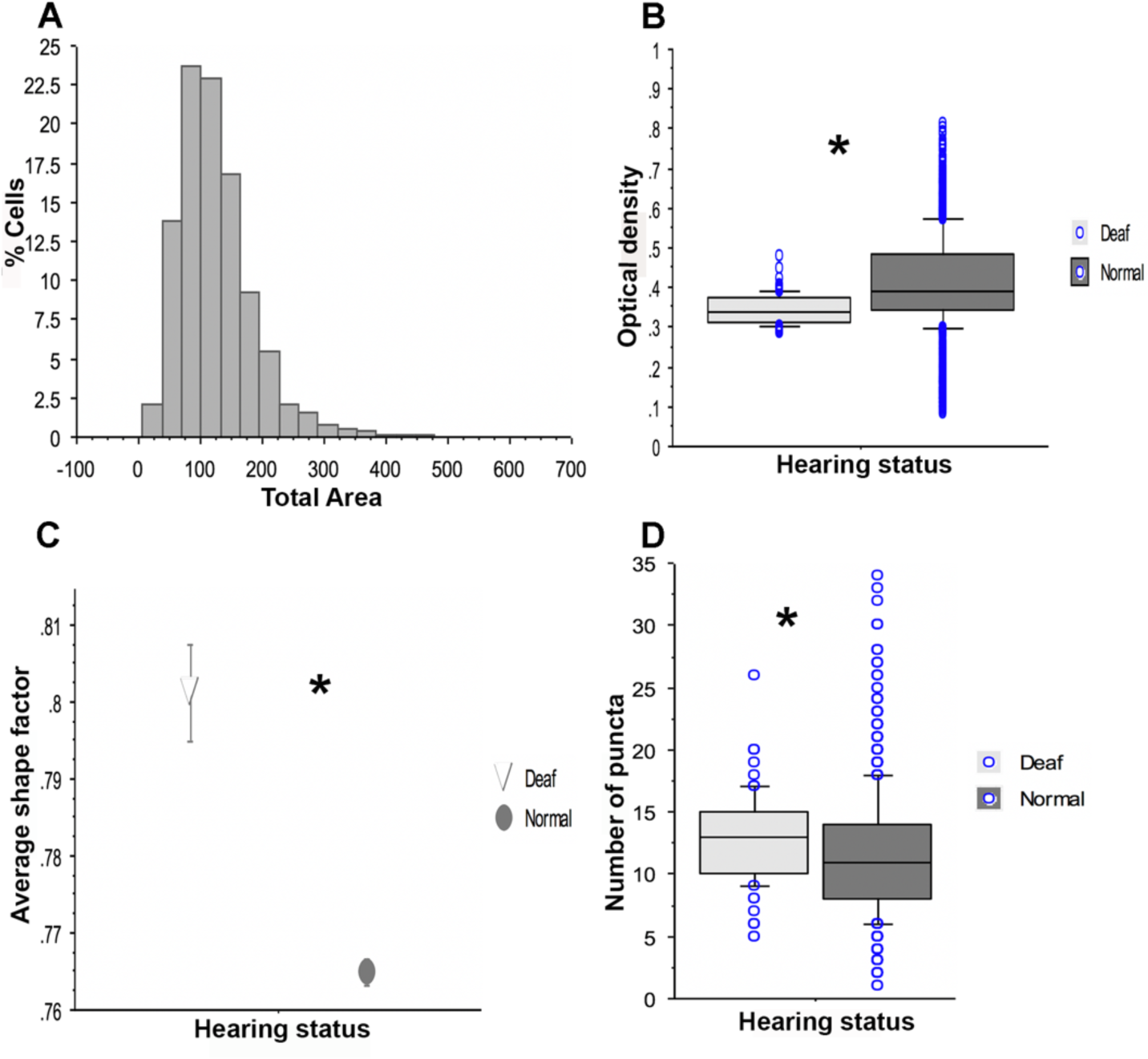
The effect of deafening on morphological features of inferior colliculus neurons. The area of somata (A), optical density (B), shape factor (C) and the number of GABA immunolabeled puncta were assessed in the central nucleus of the inferior colliculus of rats with normal hearing and compared to those deafened for 14 days. Circles are the range of values making up the sample sizes (B, D); Asterisks (*) denote statistical significance (*p* ≤ 0.05 is significant)

#### Axosomatic

The number of axosomatic puncta were counted in seven sections (28 regions encompassing 14 high and 14 low frequency regions in each subject) for a total of 4057 somata. In the normal hearing group, there was no significant difference in the number of GABA immunoreactive puncta found on somata in high frequency regions (11.85 ± 0.154) when compared to low frequency regions (11.74 ± 0.150). However, there were significantly more GABA positive puncta found apposing unlabeled cells (11.50 ± .219) when compared to GABA labeled cells (10.27 ± 0.458; p<0.045; Table 1). Administration of neomycin resulted in profound hearing loss (no detectable ABR responses at 100 dB SPL) in each of the animals. Following fourteen days of deafness the average optical density of somata in the deafened group was significantly decreased by 16.3% (Figure 4B). In addition, the number of GABA immunoreactive puncta in close apposition to somata was also modestly, but significantly, increased (Figure 4D). There were not significant changes across tonotopic regions.

**Table 1.** Number of somatic puncta in the CNIC as a function of frequency (high or low) and intensity of GABA labeling (labeled versus unlabeled). * = *p* ≤ 0.05 is significant

| High Frequency | Low Frequency | Unlabeled | Labeled | High Frequency Unlabeled | Low Frequency Unlabeled | High Frequency Labeled | Low Frequency Labeled |
| --- | --- | --- | --- | --- | --- | --- | --- |
| 11.85 ± 0.154 | 11.74 ± 0.150 | *11.50 ± 0.219 | *10.27 ± 0.458 | 11.87 ± 0.273 | 12.38 ± 0.307 | 12.11 ± 0.254 | 11.23 ± 0.280 |

#### Axodendritic

To determine whether these qualitative changes were indicative of changes in GABAergic labeling of axodendritic puncta, an assessment of the percent area labeled and optical density of non-somatic labeling, excluding the somata and puncta apposing somata, was made by thresholding each image that had been previously analyzed and comparing the threshold levels from the deafened group with those of the normal hearing group. There was a significant (26.53%) decrease in the percent thresholded area and decrease in the number of puncta in the neuropil of the deafened group when compared to the non-deafened group (Table 2, Figure 3). The results indicate that the deafened group had less GABA staining than the normal hearing group and that the primary difference may lie in the labeling of axodendritic puncta. However, we do not know whether the observed decrease in GABA labeling is due to a decrease in or a loss of GABA production or because there is a reduction in the number of puncta within the CIC following deafness. To address these questions, an ultrastructural analysis of changes with deafness in the CIC was undertaken.

**Table 2.** GABA labeled puncta in the neuropil of the CNIC as a function of frequency (high or low) and hearing status (deaf or normal). * = *p* ≤ 0.05 is significant (normal versus deaf)

| High Frequency | Low Frequency | Normal | Deaf | High Frequency Normal | High Frequency Deaf | Low Frequency Normal | Low Frequency Deaf |
| --- | --- | --- | --- | --- | --- | --- | --- |
| 13.66 ± 2.17 | 13.15 ± 1.63 | 14.51 ± 1.73 | 10.66 ± 0.122 | 14.91 ± 0.292 | 10.55 ± 0.009 | 14.11 ± 2.18 | 10.77 ± 0.253 |

### Electron Microscopy

#### General morphology

Comparisons across axodendritic (Figure 5, 6) and axosomatic (Figure 7, 8) boutons were assessed using post-embedding immunogold labeling for GABA at the ultrastructural level in hearing and neomycin administered groups (Figure 5 and 7, 6 and 8). Ultrastructural analysis reveals the same range of neuronal shapes observed at the light level. Neurons of the inferior colliculus contain mitochondria that range from small and round to hyper-elongated. The cytoplasm is filled with rough endoplasmic reticulum and studded with free ribosomes. A variety of boutons was observed in the IC containing either round, pleomorphic, or flat vesicles. Many boutons were also found to contain dense core vesicles (arrow heads in Figures 5, 7).

**Figure 5.**
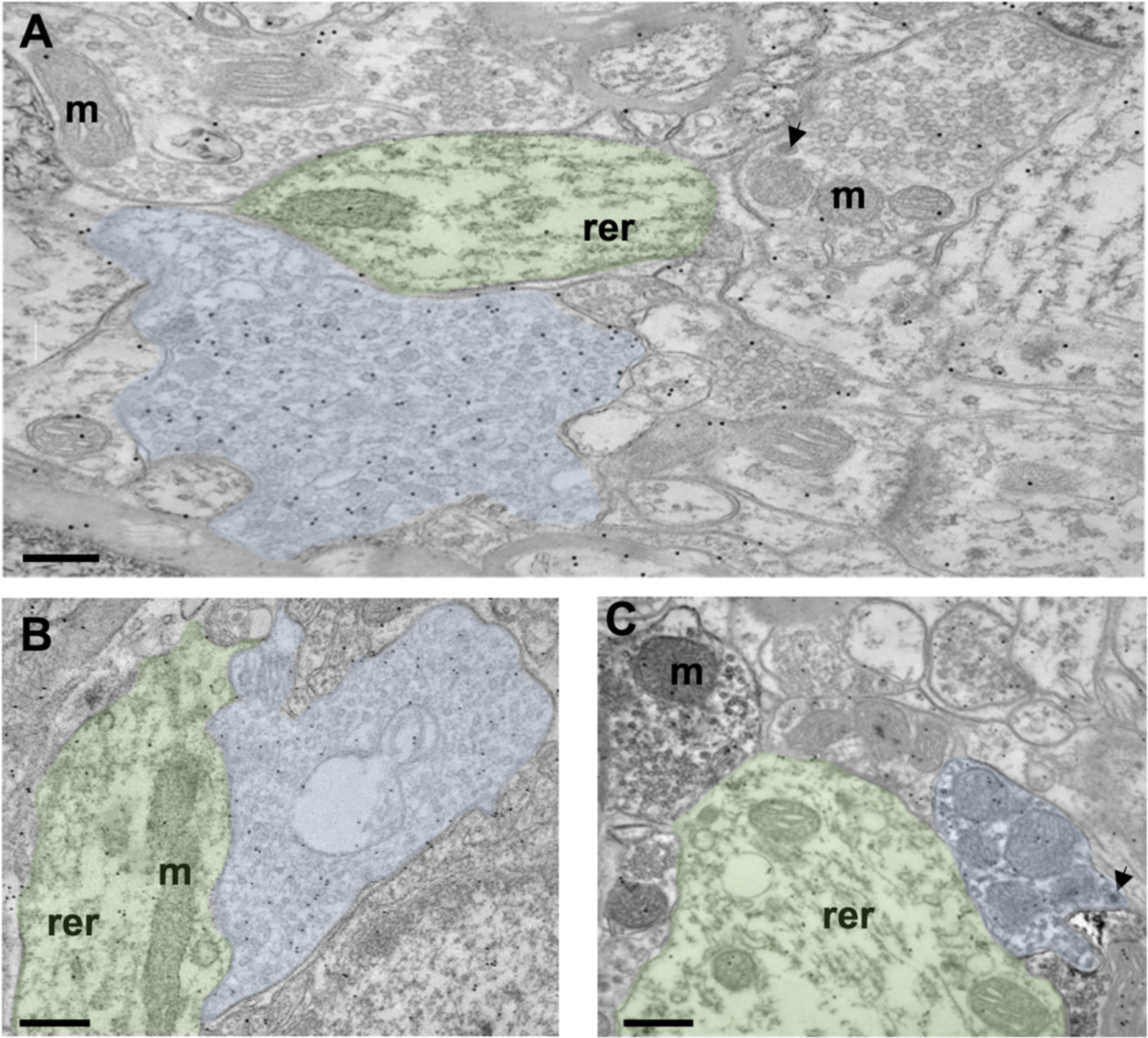
Immunogold labeling of GABAergic boutons in the inferior colliculus of normal hearing rats. Labeling for GABA is found in axodendritic boutons (blue) and in GABA immunoreactive dendrites (green; A, B, C). In normal hearing animals, robust GABA labeling is seen in boutons containing pleomorphic vesicles (A, B) while fewer immunogold particles are found in boutons containing round vesicles (A, C). m - mitochondria; rer - rough endoplasmic reticulum. Arrowheads (A,C) indicate dense core vesicles. Scale bar = 200µm

**Figure 6.**
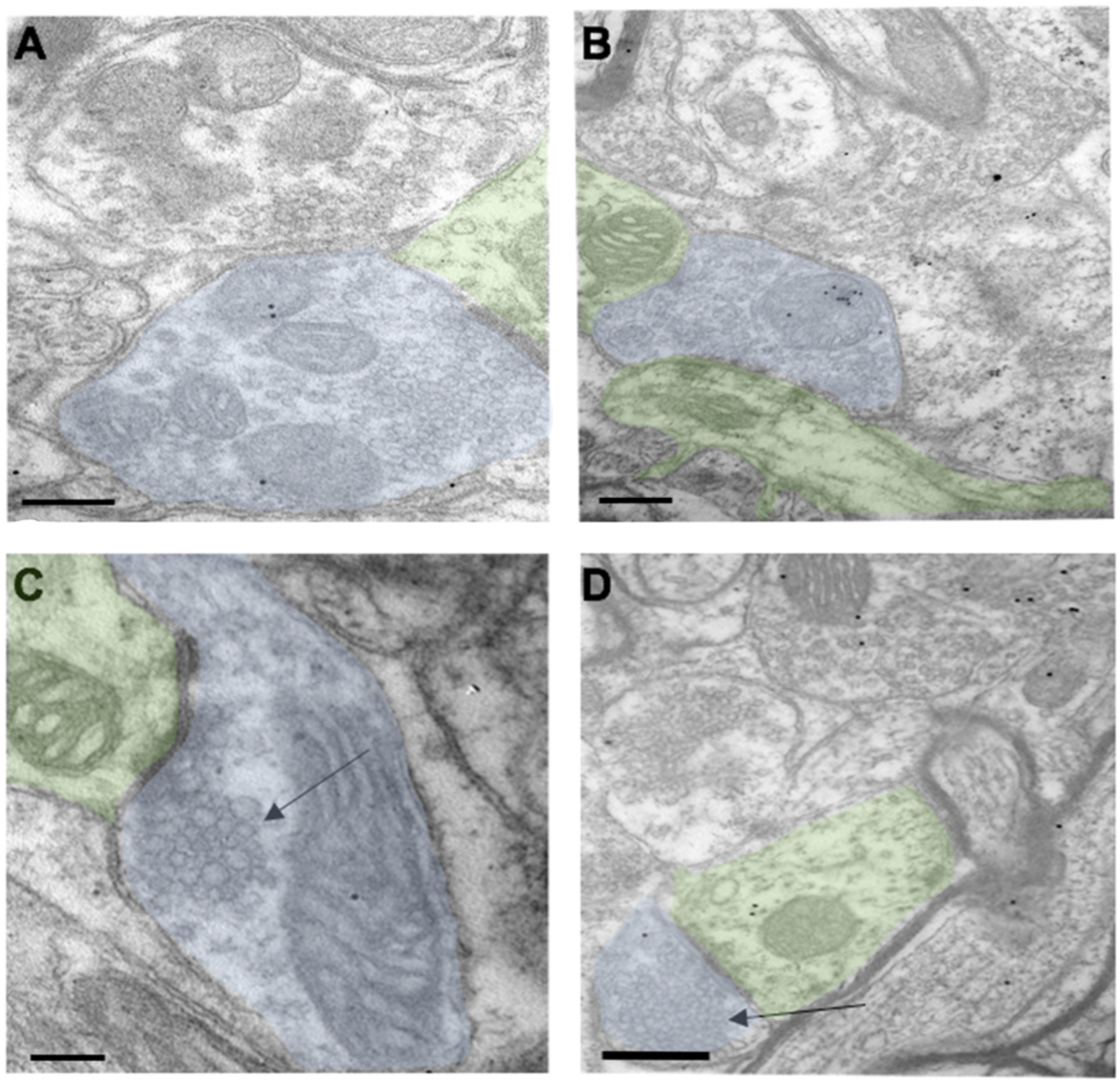
Axodendritic immunolabeling for GABA in the inferior colliculus of a deaf rat. Dendrites (green) are in synaptic contact with boutons (blue) containing pleomorphic vesicles (A, B), or round vesicles (arrows in C, D). Scale bar = 200µm

**Figure 7.**
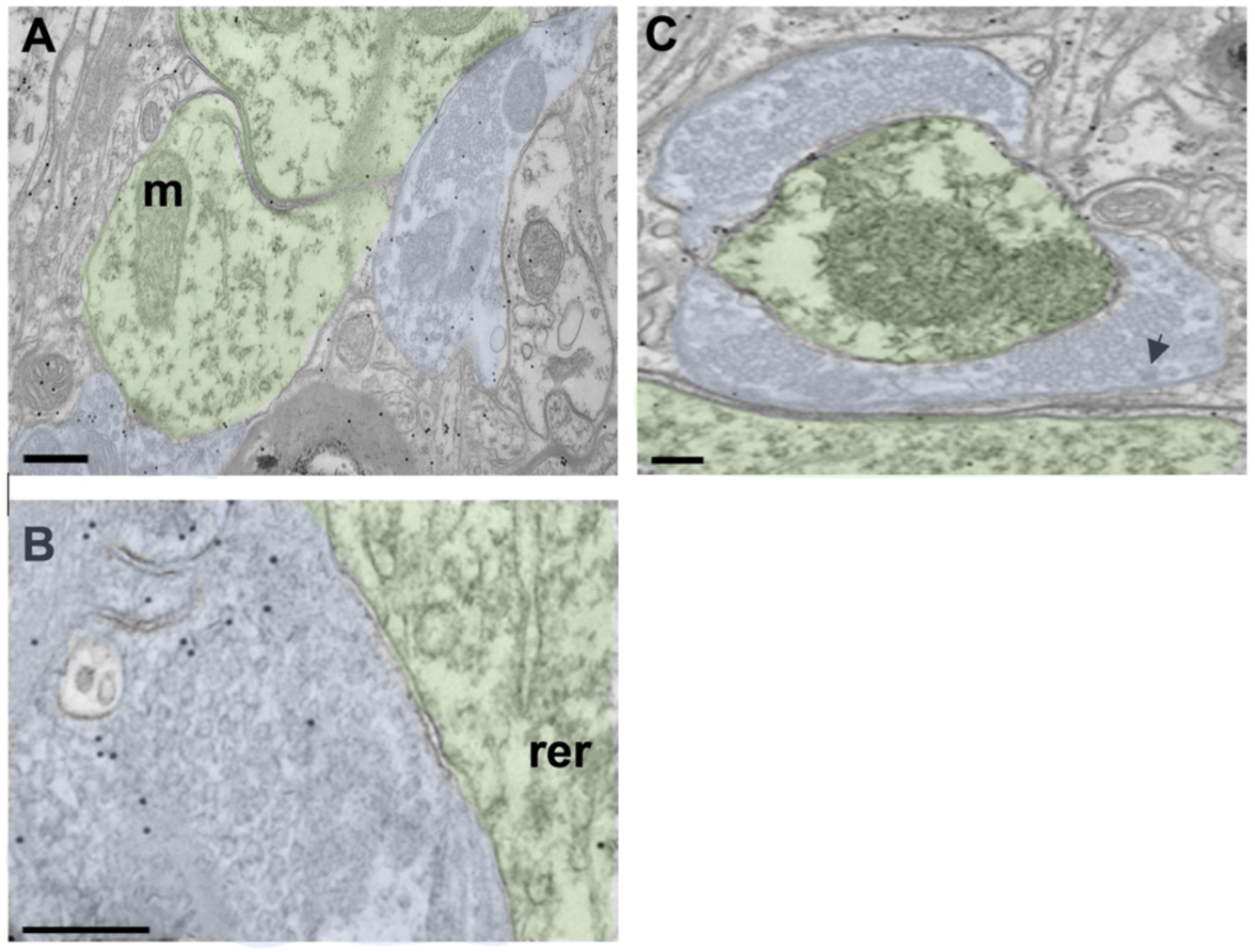
Axosomatic immunolabeling for GABA in the inferior colliculus of normal hearing rats. Labeling for GABA is found in axosomatic boutons (blue) and somata (green). Immunogold labeling for GABA is found in boutons containing pleomorphic vesicles (A, B) with few immunogold particles are located in boutons with small round (C). m - mitochondria; rer - rough endoplasmic reticulum; arrowhead - dense core vesicle; Scale bar = 200µm

#### Axosomatic / Axodendritic

Immunoreactivity for GABA was observed throughout the cytoplasm within somata, dendrites and puncta. Often the gold particles were associated with mitochondria. GABAergic immunogold labeling in axosomatic boutons was examined in normal and deafened animals. In normal hearing animals, robust GABA labeling was observed in axosomatic boutons containing pleomorphic vesicles (Figure 7). In deafened animals (Figure 8) the number of gold particles within axosomatic boutons decreased significantly (78%; p=0.0078 Table 3). The changes observed in axodendritic boutons were also dramatic (Figure 5, 6, Table 3) with the number of gold particles in boutons containing pleomorphic vesicles decreasing by 82% (p= 0.0210).

**Figure 8.**
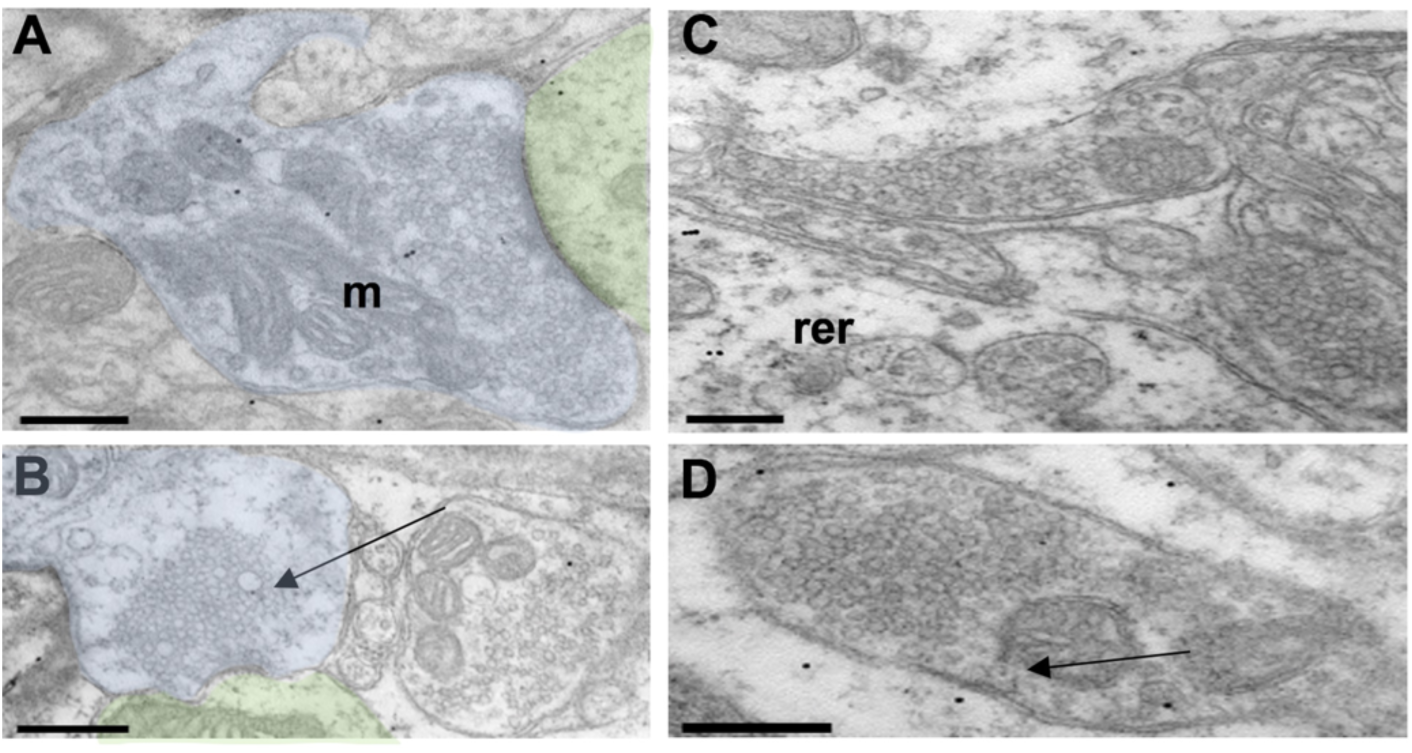
Axosomatic immunolabeling for GABA in the inferior colliculus of a deaf rat. Somata (green), can appose boutons (examples are depicted in blue A-B) containing pleomorphic vesicles (A) that are sparsely labeled or boutons containing round vesicles (arrow) with fewer immunogold particle (B). Scale bar = 200µm

**Table 3.**
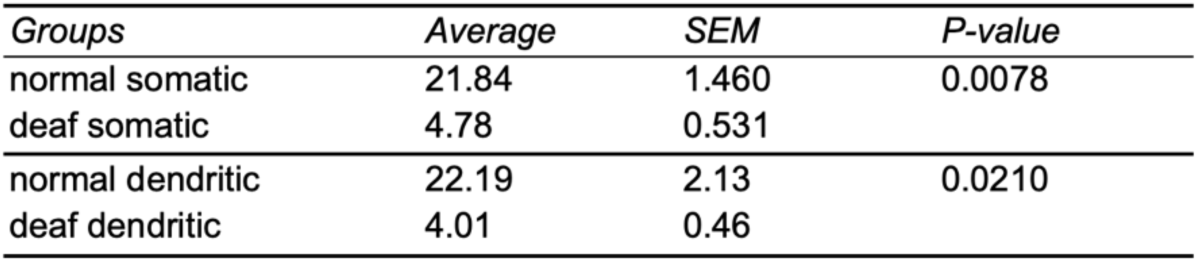
Comparison of the number of gold particles in normal and deaf axosomatic puncta as well as those found in normal and deaf axodendritic puncta. *p* ≤ 0.05 is significant

## Discussion

Robust labeling of GABAergic cells, fibers and puncta was evident in normal hearing animals. Our assessment characterizing neurons and puncta from normal subjects generally agree with other studies examining GABA immunolabeling in the CNIC, with GABAergic neurons comprising approximately 27% of all neurons in this study. Following acute bilateral deafness generated by ototoxicity using the antibiotic, neomycin, the present study demonstrates a significant decrease in GABA immunolabeling in the central nucleus of the inferior colliculus (CNIC). This finding aligns with previous research on age-related changes in the auditory system. Caspary et al. (1995) reported multiple GABA-related alterations in the aging IC, including decreased numbers of GABA immunoreactive neurons, reduced GABA release, and changes in GABA receptor binding. These age-related changes suggest altered GABAergic function in the IC, which may contribute to central auditory processing deficits associated with presbycusis.

The distribution of large and small GABA neurons in the CNIC match well with what has been reported previously (Ito et al., 2009). We also confirmed that GABAergic neurons were larger than unlabeled neurons in the CNIC (Altschuler et al., 2008; Riquelme,et al., 2001; Merchan et al., 2005; Schofield 2018; Beebe et al., 2018; Ito et al., 2018). Similar to other studies unlabeled neurons have more axosomatic GABA labeled puncta when compared to GABAergic neurons (Merchàn et al., 2005; Beebe et al., 2016). One study suggested that inhibition is decreased within each tonotopic layer, with the center having the most inhibition and the periphery showing the least (Hage and Ehret, 2003). In the current study, differences in size and number of puncta were not related to tonotopic frequency regions. This correlates well with Merchàn et al., (2005) which found similar differences to be minimally related to frequency regions within the CNIC. Interestingly, assessment of shape factor and optical density in deafened animals revealed that CNIC neurons were significantly rounder and had less labeling for GABA when compared to normal hearing animals. The current study’s observations of morphological and biochemical changes in GABAergic neurons following deafness imply potential deficits in the integration of ascending and descending signaling. This is particularly relevant given the extensive dendritic branching of large GABAergic neurons in the CNIC, which are capable of receiving and integrating multiple ascending and descending inputs to the CNIC (Oliver et al., 1991; Saldaña et al., 1996; Beebe et al., 2016, Bajo et al., 2007).

The observed reduction in GABA labeling intensity and the number of GABAergic puncta in the neuropil suggests a decrease in dendritic inhibition within the CNIC. In normal hearing animals, GABA immunogold labeling at the ultrastructural level was predominantly in terminals containing oval/pleomorphic vesicles, while terminals with large round, small round, or flattened vesicles were less labeled. Fixation artifact offers the opportunity to associate the shape of vesicles within boutons with the amino acid neurotransmitter contained within the vesicle (reviewed in Roberts and Ribak 1987). Round vesicles are associated with excitatory neurotransmitters such as glutamate and aspartate. Puncta labeled for vGluT1 and vGlut2, markers for excitatory terminals, have been reported to primarily be present in primarily distinct populations of terminals within the CNIC (Altschuler et al., 2008; Fyk-Kolodziej et al., 2011). Boutons with round vesicles were in synaptic contact with both labeled and unlabeled CNIC neurons. Flattened vesicles are linked with the inhibitory neurotransmitter glycine and pleomorphic vesicles are associated with the neurotransmitter GABA. The ultrastructural analysis in the present study revealed a significant decrease in GABA-positive immunogold particles within axodendritic and axosomatic terminals containing oval/pleomorphic vesicles following deafness. This reduction in GABA levels within IC terminals supports the hypothesis that GABAergic changes contribute to the altered balance between inhibition and excitation observed in the deafened auditory system. Such changes in inhibitory neurotransmission may underlie the neuronal hyperactivity associated with various forms of hearing loss, including noise-induced hearing loss and tinnitus (Turecek et al., 2023).

The reorganization of GABAergic microcircuits observed in this study and others suggests a common mechanism of sensory system plasticity across modalities. In the visual cortex, dark rearing has been shown to prevent the normal developmental increase in GABAergic function (Griffen et al., 2014). Additionally, Keck et al. (2011) demonstrated that in the adult mouse visual cortex, loss of sensory input leads to rapid and lasting reductions in the number of inhibitory cell spines and boutons. Similarly, in the somatosensory system, whisker removal in young rodents leads to decreased numbers of GABA-immunoreactive neurons and synaptic inputs in the corresponding cortical areas (Miraucourt et al., 2012).

These findings collectively underscore the importance of GABAergic plasticity in sensory system adaptation to altered inputs. The rapid structural changes observed in inhibitory neurons across sensory modalities suggest that GABAergic circuit reorganization may be a fundamental mechanism for maintaining the balance of excitation and inhibition in sensory processing. Future research should focus on elucidating the molecular mechanisms underlying these structural changes and exploring potential therapeutic interventions to modulate GABAergic function in sensory disorders.

In conclusion, the present study’s findings of GABA-related changes in the CNIC following deafness, when considered alongside similar observations in aging and other sensory systems, highlight the critical role of inhibitory circuit plasticity in sensory processing and adaptation. Understanding these mechanisms may provide valuable insights for developing treatments across sensory disorders, including age-related hearing loss and other forms of sensory dysfunction.

## Acknowledgements

These studies were supported by NIDCD Grant DC00383, NIDCD, NIH core center grant P30 DC05188, Department of Veterans Affairs RR&D Merit Award grant 1I01 RX001095 to AGH and American Tinnitus Association grant to AGH.

